# Extracellular matrix deposition precedes muscle-tendon integration during murine forelimb morphogenesis

**DOI:** 10.1101/2022.01.23.477427

**Authors:** Yue Leng, Sarah N. Lipp, Ye Bu, Hannah Larson, Kathryn R. Jacobson, Sarah Calve

**Affiliations:** Weldon School of Biomedical Engineering, Purdue University, 206 South Martin Jischke Drive, West Lafayette, IN 47907; The Indiana University Medical Scientist/Engineer Training Program, Indianapolis, IN 46202; Paul M. Rady Department of Mechanical Engineering, University of Colorado – Boulder, 1111 Engineering Dr, Boulder, CO 80309; Purdue University Interdisciplinary Life Science Program, 155 S. Grant Street, West Lafayette, IN 47907

**Keywords:** myotendinous junction, extracellular matrix, *Pax3*, musculoskeletal development, 3D imaging

## Abstract

The development of a functional vertebrate musculoskeletal system requires the combination of contractile muscle and extracellular matrix (ECM)-rich tendons that transmit muscle-generated force to bone. Despite the different embryologic origins, muscle and tendon integrate at the myotendinous junction (MTJ) to seamlessly connect cells and ECM across this interface. While the cell-cell signaling factors that direct development have received considerable attention, how and when the ECM linking these tissues is deposited remains unknown. To address this gap, we analyzed the 3D distribution of different ECM and the influence of skeletal muscle in forelimbs from wildype (WT) and muscle-less *Pax3*^*Cre/Cre*^ mice. At E11.5, prior to MTJ integration, an aligned ECM was present at the presumptive insertion of the long triceps into the WT ulna. Mechanically robust tendon-like and muscle compartmentalization structures, positive for type I collagen, type V collagen, and fibrillin-2, still formed when muscle was knocked out. However, MTJ-specific ECM was not observed when muscle was absent. Our results show that an ECM-based template forms independent of muscle, but muscle is needed for the proper assembly of ECM at the MTJ.

**Summary statement:** An aligned ECM template connects tendon and muscle during limb development, independent of muscle progenitor migration into the limb; however, the assembly of MTJ-specific ECM requires the presence of muscle.

## Introduction

The extracellular matrix (ECM) is a three-dimensional (3D) network that provides cells with biochemical and structural cues (Tonti et al., 2021). In adult skeletal muscle, the ECM is organized into three layers, the endomysium, perimysium, and epimysium. Each layer has a different ECM composition, ranging from the basement membrane-rich endomysium to the fibrillar collagen-rich epimysium (Muhl et al., 2020). The epimysium is thought to be continuous with the tendon to facilitate the transmission of muscle-generated contractile forces to bone for motion (Purslow, 2020; Turrina et al., 2013). Tendons are predominantly comprised of type I collagen fibers oriented along the loading direction, providing tensile strength, with a distinct pericellular ECM that surrounds linear arrays of tenocytes (Jacobson et al., 2020a; Ritty et al., 2003). Linking these tissues is the myotendinous junction (MTJ), a specialized interface marked by a localized enrichment of ECM, including type XXII collagen (COL22A1) and periostin (POSTN) (Jacobson et al., 2020a). While the structure and composition of adult muscle, tendon, and MTJ ECM are well described, how the ECM components of these disparate tissues are deposited and integrated during development is less clear.

The cells of the musculoskeletal system have different embryological origins, yet seamlessly integrate during forelimb morphogenesis (Nassari et al., 2017). In the mouse, the limb bud develops at embryonic day (E)9.5 as an outgrowth of the lateral plate mesoderm and includes tenocyte precursors (Schweitzer et al., 2001a). Scleraxis (*Scx*), a transcription factor expressed in developing tendons, is detected in limb buds at E10 (Schweitzer et al., 2001b). Muscle progenitors migrate into the limb from the somite starting around E10, which requires the transcription factor Paired box gene 3 (*Pax3)* (Bober et al., 1994). Muscle precursors then differentiate and fuse into myofibers (Deries and Thorsteinsdóttir, 2016). By E12.5, tendon progenitors are aligned between muscle and cartilage, and the basic pattern is complete by E13.5 (Hasson, 2011; Murchison et al., 2007). MTJ development is hypothesized to occur simultaneously with the condensation of tendon and muscle progenitors (Narayanan and Calve, 2021; Subramanian and Schilling, 2015); however, fibrillar collagen deposition is not thought to take place until after the progenitors reach their final destination, reportedly starting around E14.5 (Revell et al., 2021).

Notably, interactions between tendon and muscle are not required for initial musculoskeletal patterning. Tendons form in the appropriate position in muscle-less limbs (Huang et al., 2015; Kardon, 1998; Kieny and Chevallier, 1979), and muscle cells aggregate in the correct location even when tendon development is disrupted (Pryce et al., 2009), suggesting patterning and subsequent integration is regulated by factors autonomous of muscle and tendon progenitors, such as connective tissue cells and the ECM.

Perturbation of transcription factors that regulate connective tissue cell behavior (*e*.*g. Tbx3, Tbx5, Osr1, Osr2*) disrupt musculoskeletal patterning, and it has been hypothesized that the ECM plays an instructive role in muscle-tendon integration (Helmbacher and Stricker, 2020; Shellswell et al., 1980). However, the distribution of the ECM and how it relates to muscle-tendon development remains poorly defined. To determine the spatio-temporal distribution of the ECM during limb development, and the influence of skeletal muscle on matrix deposition, we used tissue clearing, 3D imaging, and proteomics to investigate wildtype (WT) and muscle-less *Pax3*^*Cre/Cre*^ limbs. Our data revealed that an ECM-based template forms between these tissues, independent of muscle cell migration and MTJ formation.

## Results and Discussion

### ECM alignment preceded muscle-tendon patterning

To investigate the organization of ECM fibers prior to the establishment of the final muscle and tendon pattern, WT murine forelimbs were analyzed at E11.5. The resolution of the ECM in 3D was enhanced by decellularization or clearing using SeeDB (Figure 1A-D’; see Methods). In decellularized E11.5 forelimbs, type I collagen (COL1) and WGA^+^ proteoglycans were assembled in a linear, fibrillar structure that emanated from the ulna where the presumptive long triceps will insert (Figure 1E-E’’’). To visualize the distribution of myogenic progenitors in the context of the ECM, *Pax3*^*Cre*^*/ZsGreen1*^*+*^ (*Pax3*GFP^+^) limbs were cleared using SeeDB. These tissues were also counterstained for ECM highly expressed in the developing limb: tenascin-C (TNC), fibrillin-2 (FBN2), and type V collagen (COL5) (Jacobson et al., 2020b). Proximally, *Pax3*GFP^+^ cells were loosely aggregated (Figure 1F-H’’; *), whereas they were oriented along the proximal-distal (p – d) axis of the limb towards the distal tip of the limb (arrowhead). FBN2^+^ and COL5^+^ fibers were also aligned along the p – d axis, extending distally ahead of the migrating *Pax3*GFP^+^ cells (Figure 1G’-H’’; arrowhead).

**Figure 1:**
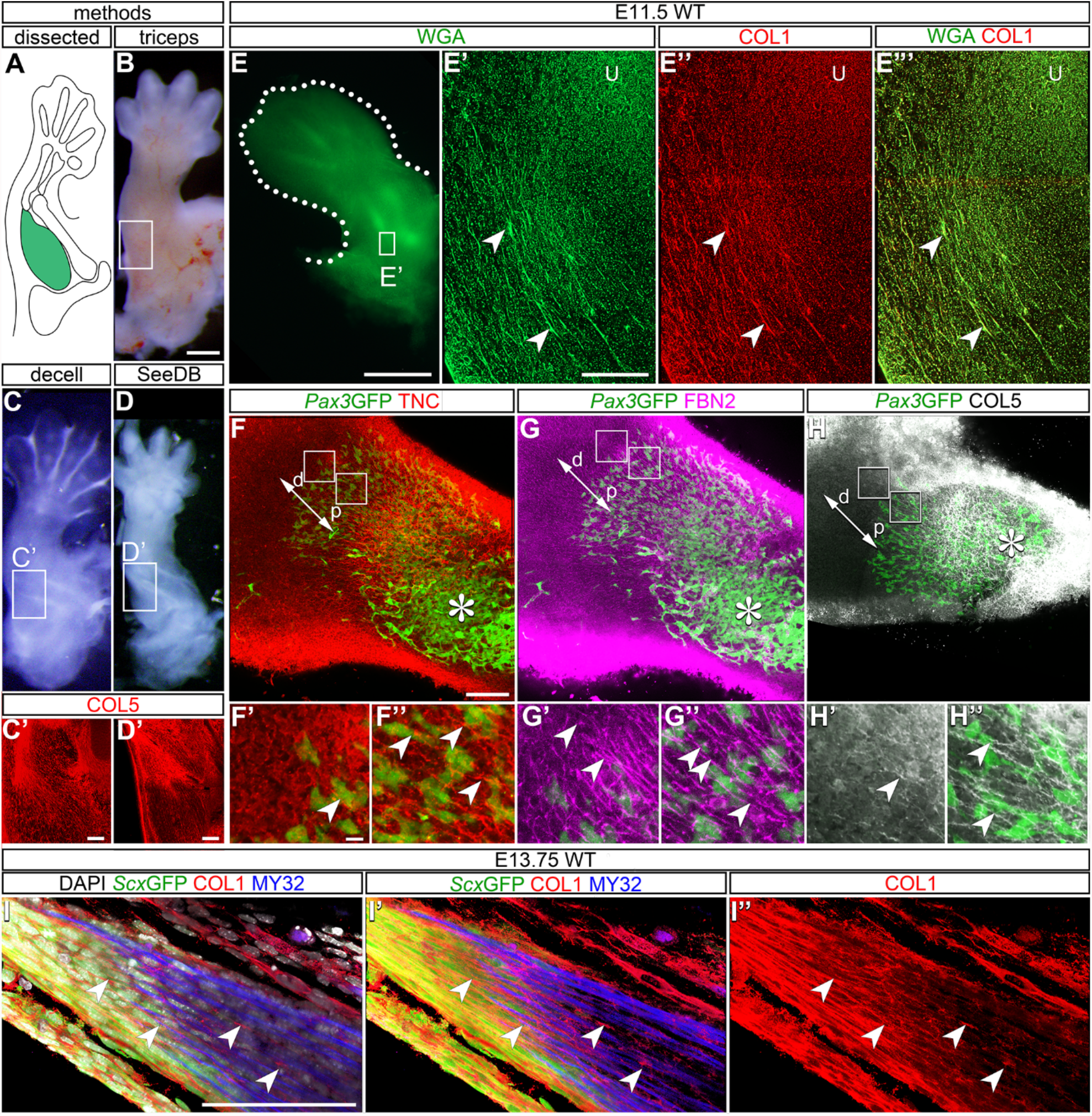
ECM alignment preceded muscle-tendon patterning. **(A-D’)** To visualize the 3D ECM structure of the triceps (green, A), murine forelimbs (B) were decellularized (decell) in 0.05% sodium dodecyl sulfate (SDS; C) or optically cleared using SeeDB (D). Samples were imaged using confocal microscopy and maintained consistent organization of the ECM (COL5; red) after both decell (C’) and SeeDB (D’). Representative images from E13.5-E13.75 forelimbs. **(E-E”’)** Aligned COL1^+^ (red) and WGA^+^ fibers (green) inserted into the lateral triceps near the ulna (U) in decellularized WT E11.5 forelimbs. **(F-H)** In SeeDB-cleared E11.5 forelimbs, *Pax3*GFP^*+*^ cells (green, *) aligned along the proximal (p) - distal (d) axis (arrow) with TNC (red), FBN2 (magenta), and COL5 (grey). Insets show aligned ECM networks distal to *Pax3*GFP^*+*^ cells (G’,H’) and around *Pax3*GFP^*+*^ cells (F”-H”) (arrowheads). **(I-I’’)** COL1^+^ fibers (red, arrowheads) extended between *Scx*GFP^+^ tendon (green) and My32^+^ muscle (blue) in a wholemount E13.75 zeugopod. Magnification 10× (C’), 20× (E - E’’), 25× (D’, F - H), 63× (I - I’’), z projection: z = 40 µm (B’, C’) and 11.4 µm (F - H), 3D rendering: z =12.7 µm, scale bars = 1 mm (B - E); 100 µm (C’- I’’).

Since myogenic progenitors align prior to differentiating into myofibers (Besse et al., 2020) (Figure 1F’-H”; arrowhead), the ECM may enhance alignment and provide a path to guide muscle cell migration. For example, previous studies demonstrated aligned topographical cues (Lam et al., 2006), as well as signals from specific ECM proteins (*e*.*g*. TNC, fibronectin, laminins), promoted muscle migration and differentiation (Calve et al., 2010; Silva Garcia et al., 2019; Vaz et al., 2012). The initial aligned networks may provide a template along which the ECM that ultimately links muscle and tendon is deposited; aligned COL1^+^ fibers were even more prevalent after tendon and muscle integration at E13.75 (Figure 1I-I’’; arrowhead).

### Limb ECM composition was maintained despite the absence of muscle

To investigate the contribution of muscle progenitors in depositing the ECM, the proteomes of WT and muscle-less (*Pax3*^*Cre/Cre*^) forelimbs were compared. *Pax3* is functionally knocked out in mice homozygous for *Pax3*^*Cre*^, preventing the migration of muscle progenitors into the forelimb (Engleka et al., 2005) (Figure 2A). The ECM composition was determined by isolating ECM proteins, or matrisome (Naba et al., 2016), from E13.5 limbs and analyzed via liquid chromatography-tandem mass spectrometry (LC-MS/MS) (Jacobson et al., 2020b).

**Figure 2:**
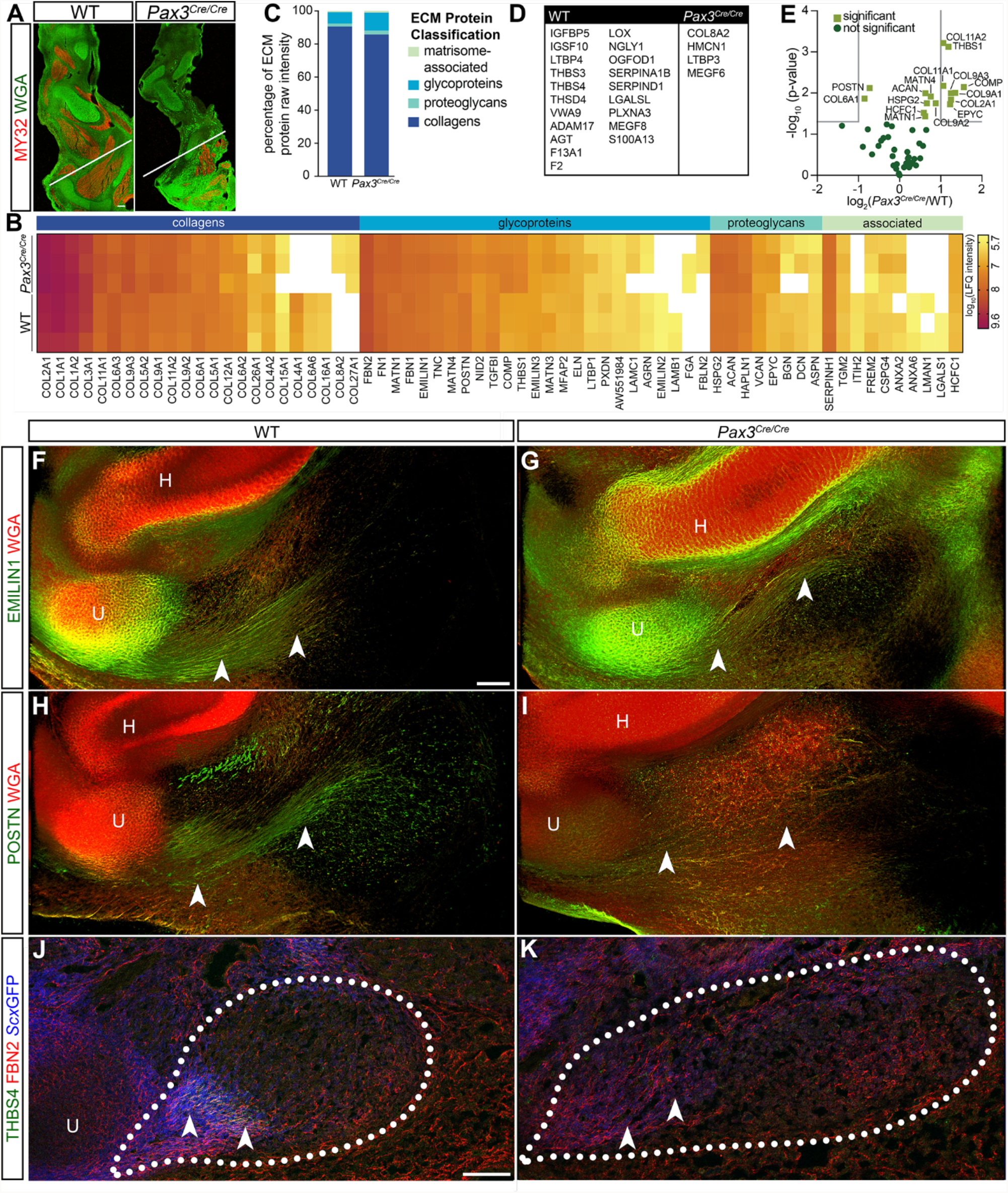
Forelimb ECM composition was minimally disrupted by the absence of muscle. **(A)** *Pax3*^*Cre/Cre*^ forelimb lacked limb musculature (MY32, myosin; red) at E13.5. The shoulder girdle begins below the line. **(B)** Log_10_(LFQ) heat map of matrisome components manually grouped via ECM protein classification. Tissues were fractionated following (Jacobson et al., 2020a), and only proteins identified in the insoluble fraction and n ≥ 2 biological replicates were included. White boxes signify zero intensity values. **(C)** Two-way ANOVA analysis revealed there were no significant differences in matrisome composition between genotypes at E13.5 (Table S2). **(D)** Matrisome components exclusively identified in either WT and *Pax3*^*Cre/Cre*^ mouse embryos for all tissue fractions. **(E)** Volcano plot comparison of log_2_(LFQ) values for the *Pax3*^*Cre/Cre*^ and WT E13.5 forelimbs, significance based on *p* < 0.05. **(F, G)** EMILIN1^+^ and WGA^+^ fibers (arrowheads) were observed in the tendons of both WT and *Pax3*^*Cre/Cre*^ embryos at E13.75. **(H, I)** Decellularized forelimb tendons from WT, but not *Pax3*^*Cre/Cre*^, forelimbs were POSTN^+^ (arrow). **(J, K)** FBN2 was retained in the *Scx*GFP^+^ area in *Pax3*^*Cre/Cre*^ forelimbs, whereas THBS4 was not. Dotted line delineates region of triceps muscle. Magnification 10× (A, F - K), 3D rendering z = 302 µm (F - I), cryosections (J, K), scale bars = 100 µm.

There was no significant difference in the overall abundance of ECM proteins between WT and *Pax3*^*Cre/Cre*^ as a function of matrisome classification (Naba et al., 2016) (Figure 2B-E). Many ECM components that are more abundant in adult tendon compared to muscle (*e*.*g*. COL1A1&2, COL5A1, EMILIN1, FBN2, TNC) (Jacobson et al., 2020a), were not altered in the *Pax3*^*Cre/Cre*^ limb (Figure 2B, F-G, S1). Additionally, the distribution of *Scx*GFP^+^ cells did not noticeably expand in the absence of muscle (Figure S1), consistent with the lack of overall change in the tendon-related matrisome.

In contrast, thrombospondin-4 (THBS4) and POSTN were exclusive to or significantly more abundant in WT limbs (Figure 2D-E), which was validated by IHC (Figures 2H-K, S1). POSTN is a matricellular protein that is elevated in tendon and muscle during repair and can promote tissue regeneration (Lorts et al., 2012; Wang et al., 2021). THBS4 is required for myoseptum stability and the expression is mechanically regulated in zebrafish (Subramanian and Schilling, 2014). The loss of POSTN and THBS4 in the tendon in the *Pax3*^*Cre/Cre*^ limb indicates deposition in the mouse also requires signals from muscle and/or muscle contraction (Figure S1). Another protein that was not identified in *Pax3*^*Cre/Cre*^ limbs was lysyl oxidase (LOX), an enzyme that forms crosslinks in many ECM (Figure 2D). Previous studies indicated that LOX is expressed by muscle cells (Gabay Yehezkely et al., 2020; Kutchuk et al., 2015), and our data suggest that myogenic progenitors may be the primary source of this enzyme in developing muscle.

A number of basement membrane-associated proteins were unaltered, which may be due to the endomysium not being fully mature at E13.5 (Leivo and Engvall, 1988), as well as retained expression in cartilage and blood vessels (*e*.*g*. NID2) (Figure S1). Indeed, cartilage-related proteins were significantly elevated (ACAN, COL2A1, COL9A1&3, COL11A1&2, COMP), or exclusively found (COL8A2, COL27A1), in the *Pax3*^*Cre/Cre*^ limb (Bian et al., 2020; Boot-Handford et al., 2003; Ocken et al., 2020), which can be explained by the relative increase of cartilage content in the absence of muscle (Figure 2E).

As the majority of the fibrillar ECM was retained in the *Pax3*^*Cre/Cre*^ limb (Figure 2B), this suggests muscle cells do not substantially contribute to total ECM deposition; however, expression of some matrisome components, such as POSTN, THSB4, and LOX are influenced by muscle signaling and contraction.

### Tendon and epimysial-like ECM formed in the absence of muscle

The minimal effect of muscle knockout on the matrisome (Figure 2), combined with the presence of tendons in *Pax3* mutant mice (Figure S1) (Bonnin et al., 2005; Huang et al., 2015), suggested that overall connective tissue patterning was maintained in the absence of muscle. Therefore, we next investigated how ECM patterning in E12.5-E14.5 forelimbs was affected in *Pax3*^*Cre/Cre*^ mutants. In WT forelimbs, WGA^+^ ECM fibers were enriched where the long triceps attached to the ulna and scapula at E12.5 (Figure 3A). At E13.5 and E14.5, fibers in the tendon bundles of WT forelimbs became more elongated and densely packed (Figure 3C, E). In *Pax3*^*Cre/Cre*^ forelimbs, WGA^+^ fibers had a similar distribution pattern as the WT at all timepoints, but were more loosely arranged in the tendon (Figure 3B, E, F).

**Figure 3:**
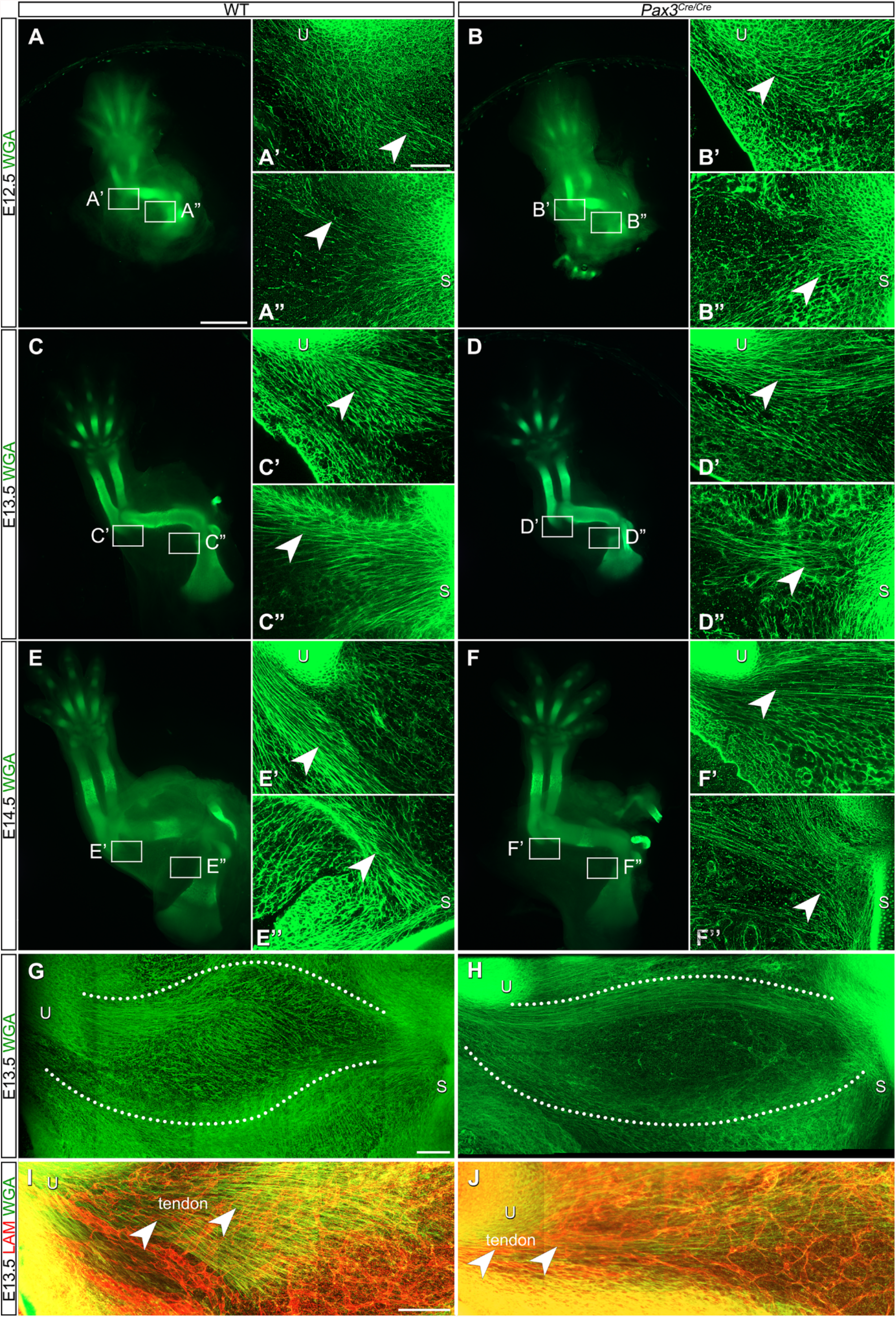
Tendon and epimysial-like ECM formed in the absence of muscle. **(A-F’’)** Decellularized E12.5-E14.5 WT and *Pax3*^*Cre/Cre*^ forelimbs were rich in WGA^+^ fibers at the insertion of the long triceps into the ulna (U; arrowhead; A’-F’) and origin of the long triceps at the scapula (S; arrowhead; A”-F”). However, the fiber bundles were less dense in *Pax3*^*Cre/Cre*^ compared to WT forelimbs. **(G, H)** For the long triceps of both E13.5 WT and *Pax3*^*Cre/Cre*^ embryos (dotted outline), WGA^+^ fibers formed epimysial-like structures. **(I, J)** In the *Pax3*^*Cre/Cre*^ forelimbs, laminin (LAM^+^) blood vessels (red) were organized similar to those in WT in the tendon (arrowhead) and muscle. Magnification 20×, z-projection: z = 230 µm (G, H), 138 µm (I), and 181 µm (J), scale bars = 1 mm (A - F), 100 µm (G - J).

Surprisingly, muscle compartmentalization by epimysial-like WGA^+^ fibril bundles (dotted line, Figure 3G, H, S2) and the 3D network of laminin (LAM^+^) blood vessels (Figure 3I, J) were conserved in *Pax3*^*Cre/Cre*^ forelimbs at E12.5-E13.5, indicating epimysial and vascular networks organized independently of muscle. Two types of ECM organization were observed regarding the tendon insertion of the long triceps into the ulna in both WT and *Pax3*^*Cre/Cre*^ forelimbs (Figure 3A-H): (1) fibers emanated from the tendon through the muscle belly and (2) epimysial-like fibers outlined the contour of the triceps muscle.

While several mechanisms could regulate muscle splitting (Hasson, 2011), our data support the hypothesis that accumulations of connective tissue cells regulate muscle patterning through ECM fiber deposition (Helmbacher and Stricker, 2020; Schroeter and Tosney, 1991). The persistence of vasculature in *Pax3*^*Cre/Cre*^ forelimbs may also contribute to patterning as platelet-derived growth factor B (PDGFB) from blood vessels mediated connective tissue cell compartmentalization by promoting secretion of ECM (Tozer et al., 2007). Nevertheless, muscle is needed for maturation of the overall ECM. The less robust ECM bundles observed in *Pax3*^*Cre/Cre*^ tendons could result from the lack of muscle-derived signals (*e*.*g*. biochemical, mechanical) that are needed for tendon maturation (Bonnin et al., 2005; Huang et al., 2015). The decrease in the number of WGA^+^ fibers within the muscle belly region may be due to the lack of muscle-derived LOX crosslinking (Figure 2) to stabilize the ECM and make it less resistant to extraction via SDS.

### Fibrillar ECM persisted in the muscle belly of *Pax3*^*Cre/Cre*^ mutants but MTJ-specific ECM did not

To further characterize the ECM in developing muscle, the composition and mechanical integrity of decellularized E13.5 WT and *Pax3*^*Cre/Cre*^ forelimbs were investigated. In the WT limb, COL1^+^, COL5^+^, and FBN2^+^ fibers were concentrated in the tendon and remained highly aligned crossing through the muscle belly. In the mutant limb, COL1 staining outlined an epimysial-like structure where the muscle should be and COL5^+^ fibers were more sparse compared to the WT limb. However, FBN2^+^ fibers were still present across the muscle belly region, similar to WT littermates (Figure 4A-F, arrowhead).

**Figure 4:**
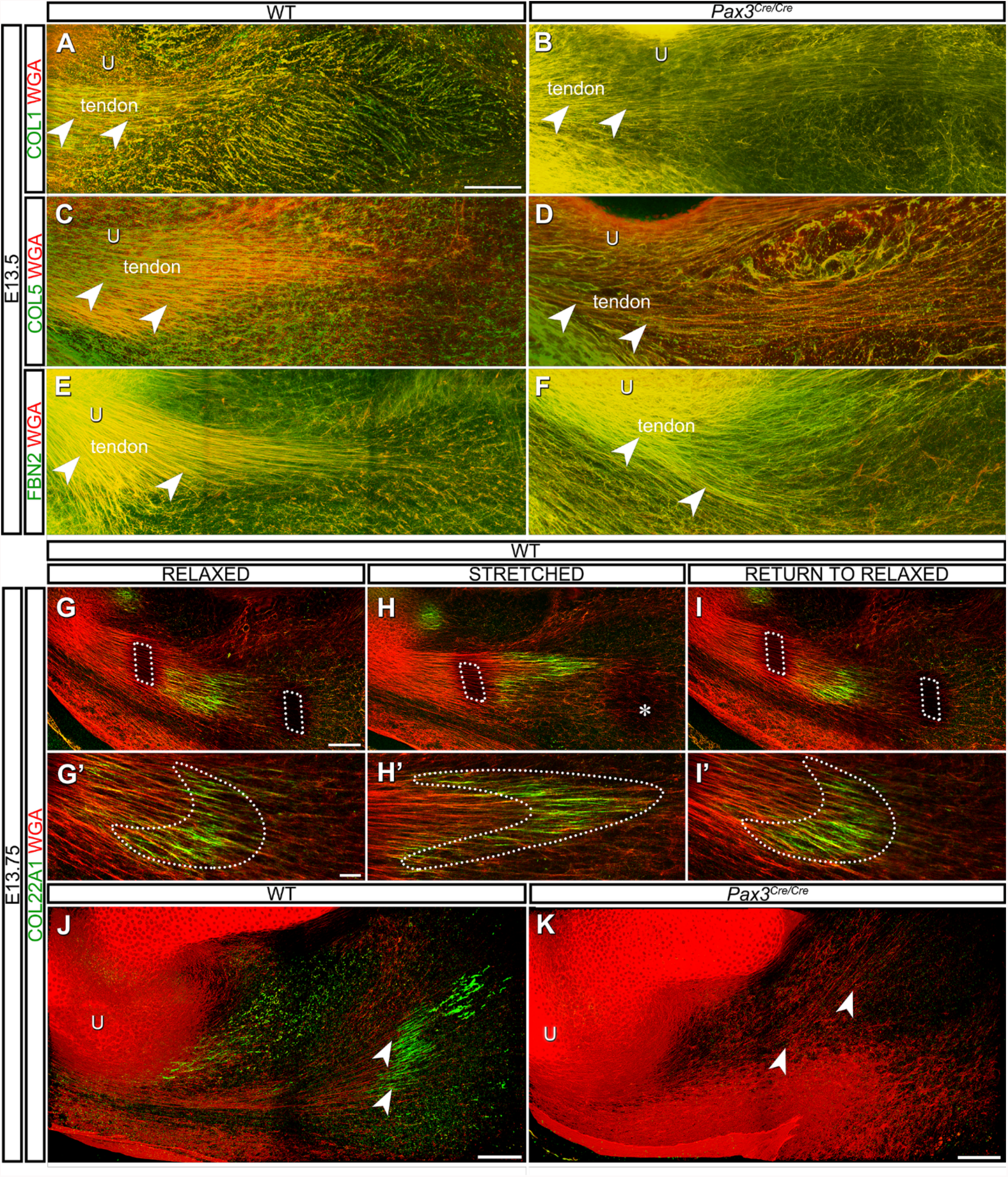
Fibrillar ECM persisted in the muscle belly of *Pax3*^*Cre/Cre*^ mutants but MTJ-specific ECM did not. **(A-F’’)** WT or *Pax3*^*Cre/Cre*^ decellularized E13.5 triceps tendon (dotted outline) and muscle belly fibers were COL1^+^, COL5^+^, and FBN2^+^. **(G-I)** In WT decellularized E13.75 forelimbs, the COL22A1^+^ MTJ (green) and WGA^+^ ECM fibers (red) were restored to original configuration when tension was removed, as indicated by the fiducial markers (dotted outline). * denotes an unresolved fiducial marker. **(G’-I’)** The MTJ exhibited elongation in under tension (dotted outline). **(J-K)** The MTJ of decellularized WT but not *Pax3*^*Cre/Cre*^ forelimbs was marked by COL22A1^+^ (green) at the end of the WGA^+^ tendon fibers (red). Magnification = 20× (A-F), 25× (G-K). Z projection: z = 230 µm (A), 122 µm (B), 24 µm, (C), 23.6 µm (D), 230 µm (E), 115 µm (F), 96.3 µm (G, G’, H, H’), 96.5 µm (I, I’), 3D rendering: z = 110 µm (J, K), scale bars = 100 µm (A-K), 25 µm (G’-I’).

The persistence of ECM in the tendon of *Pax3*^*Cre/Cre*^ mutants suggests the initial deposition of matrix by tendon progenitors is not dependent on muscle (Boregowda et al., 2008; Sengle et al., 2015). In contrast, the decrease in fibrillar collagens in the muscle belly region of *Pax3*^*Cre/Cre*^ forelimbs may be due to missing reciprocal interactions between muscle and connective tissue cells (*e*.*g*. crosslinking by LOX) needed to stabilize the fibrillar structure. Alternatively, the fibers still formed but collapsed to the periphery of the muscle belly in the absence of myofibers.

Notably, the ECM within the tendon and across the MTJ is mechanically robust. E13.75 forelimbs were decellularized, but not fixed with paraformaldehyde prior to staining for COL22A1, a matrisome component found only at the interface of muscle and tendon in the adult (Jacobson et al., 2020a). Similar to the adult, COL22A1 was only observed at the MTJ in E13.75 forelimbs. The stylopod was loaded under tension using a custom tensile tester (see Methods) (Acuna et al., 2021). After displacing the actuators by 2.5 mm, the COL22A1^+^ interface elongated. The original ECM configuration was restored when the tissue was returned to the resting position (Figure 4G-I’). We then assessed if COL22A1 was present in the *Pax3*^*Cre/Cre*^ at the tip of the WGA^+^ tendon; however, COL22A1 was absent (Figure 4J, K, S3).

COL22A1 is necessary to maintain MTJ integrity in the zebrafish, where COL22A1 knockout or knockdown leads to disrupted muscle phenotypes (Charvet et al., 2013; Malbouyres et al., 2021). Together with our results demonstrating that COL22A1 interface between muscle and tendon reversibly elongates under tensile loading, these data indicate that COL22A1 connects the ECM between muscle and tendon. While multiple cell types have been hypothesized to synthesize COL22A1 at the MTJ, including muscle (Kim et al., 2020; Koch et al., 2004; Subramanian and Schilling, 2015), tendon/connective tissue (Scott et al., 2019; Yaseen et al., 2021), or both (Gaffney et al., 2021; Petrany et al., 2020), our data revealed that COL22A1 integration requires the presence of muscle in the limb.

In summary, we provide evidence that an aligned ECM is established as early as E11.5, which becomes integrated across the MTJ as the musculoskeletal system takes on the final pattern. This network is mechanically robust by E13.75 and may provide structural as well as biochemical cues to guide the formation of the muscle-tendon interface. The general pattern of the ECM in the long triceps muscle-tendon unit was not affected when muscle progenitors failed to migrate into the limbs; however, the expression of some matrisome components (*e*.*g*. COL22A1, THBS4, LOX) were dependent on the presence of muscle. Future studies that specifically disrupt connective tissue cells will provide additional critical insight into the ECM-based mechanisms that orchestrate musculoskeletal assembly.

## Materials and Methods

### Animal models and tissue collection

All murine experiments were approved by either the Purdue University or University of Colorado Boulder Institutional Animal Care and Use Committee (PACUC or IACUC; protocol #1209000723 or #2705). PACUC and IACUC ensures that all animal programs, procedures, and facilities at the University adhere to the policies, recommendations, guidelines, and regulations of the USDA and the United States Public Health Service following the Animal Welfare Act and university Animal Welfare Assurance.

*Pax3*^*Cre*^ (#005549; (Engleka et al., 2005)) and ROSA-ZsGreen1 (*ZsGreen1*; #007906; (Mao et al., 2001)) transgenic mice were obtained from the Jackson Laboratory. *Pax3*^*Cre*^/*ZsGreen1*^*+*^ were generated to label all *Pax3*-expressing cells and their progeny with GFP^+^, which also includes a small subset of endothelial cells (Hutcheson et al., 2009). Specifically, males heterozygous for the *Pax3*^*Cre*^ transgene (*Pax3*^*Cre/+*^) were time-mated with females homozygous for *ZsGreen1*. Noon of the day a copulation plug was found was designated as E0.5. *Pax3*^*Cre/+*^ heterozygous mice were crossed to generate Pax3^Cre/Cre^ embryos, which were identified by their neural tube and neural crest defects (Engleka et al., 2005) or determined by genotyping. WT controls were defined as *Pax3*^*Cre/*+^ or *Pax3*^*+/+*^. To visualize tendon progenitors, mice containing the *ScxGFP* transgene (Pryce et al., 2007), kindly provided by Ronen Schweitzer, were time mated. E10.5-E14.5 embryos were harvested from dams euthanized via CO_2_ inhalation followed by cervical dislocation. The embryos were transferred to 1× phosphate-buffered saline (PBS) on ice. Removal of the yolk sac and amnion was performed under a dissecting microscope (DFC450, Leica Microsystems) to avoid damaging the embryos. After dissection, forelimbs were immediately either fixed in 4% paraformaldehyde (PFA; Fisher Scientific) at 4°C overnight then washed with PBS for the preparation of sample for vibratome sectioning or wholemount staining, or processed for decellularization or cryosectioning.

### Forelimb decellularization

After dissection, freshly harvested forelimbs were mounted in 1% low gelling temperature agarose (Sigma-Aldrich) in a 10 mm × 10 mm× 5 mm biopsy cryomold (Tissue-Tek). The agarose cubes containing forelimbs were submerged in 1 mL solution containing 0.05% sodium dodecyl sulfate (SDS; VWR), with 2% penicillin-streptomycin or 0.02% sodium azide (Sigma-Aldrich), in PBS, and gently rocked at room temperature (RT). The SDS solution was replaced every 24-48 hours (h) until decellularization was complete, after 3-6 days. Upon decellularization, the agarose cubes were washed in 1× PBS buffer for 1 h, fixed with 4% PFA in PBS for 1 h, then washed with PBS for 1 h again with gentle rocking at RT. The decellularized forelimbs were carefully extracted from the agarose under a dissecting microscope, and stored in 1× PBS buffer at 4°C until stained and imaged.

### Vibratome sectioning and wholemount preparation

Fixed, intact forelimbs were embedded in 3.5% low melting point agarose (Amresco) in 1× PBS and set at RT for 30 min to solidify. Agarose-embedded samples were then positioned in the appropriate plane and attached to the sample holder of a Leica VT-1000S vibratome with Loctite super glue and the chamber was filled with PBS to maintain sample hydration. Forelimbs were sliced into 250 μm thick sections and placed in a tissue culture dish filled with PBS on ice. Extra agarose was dissected away from the tissue by forceps. Samples were stained via immunohistochemistry the same day.

For wholemount preparation, fixed, intact forelimbs were prepared by removing skin or embedded in 3% low melting point agarose and bisected in the dorsal-ventral plane using a scalpel.

### Cryosectioning and immunohistochemistry

Forelimbs from *Pax3*^*Cre/Cre*^ and WT embryos were fixed in 4% PFA for 1 h, washed with PBS 3 × 30 minutes (min) before embedding in Optimal Cutting Temperature compound (OCT; Sakura Finetek), frozen with dry ice-cooled isopentane (Fisher Scientific), and stored at −80 °C. Cryosections (10 µm thickness) were collected on His-bond glass slides (VWR) and processed following (Jacobson et al., 2020a). Sections were incubated with primary antibodies (Table S1) at 4°C overnight, and washed with PBS for 3 × 5 min. Slides were then stained with secondary antibodies and probes (Table S1). Sections were imaged at 10× water using a Zeiss LSM 880 confocal.

### Fluorescent labeling of ECM and imaging

After PFA fixation, decellularized forelimbs or vibratomed sections were incubated in blocking buffer [10% donkey serum diluted in 1× PBS with 1% Triton X-100 (PBST) and 0.02% sodium azide] for 16 h at 4°C to increase the ability of antibodies to permeate through the sample and to block non-specific binding. Samples were then incubated with primary antibodies (Table S1) diluted in blocking buffer, and gently rocked at 4°C for 48 h. Samples were rinsed 3 × 30 min with 1% PBST at 25°C, and then incubated with secondary staining reagents diluted in blocking buffer, placed in a lightproof container, and rocked again at 4°C for 48 h. WGA was used as a counter stain as it binds to sialic acid and N-acetylglucosamine and marks proteoglycans in the musculoskeletal architecture (Kostrominova, 2011). Finally, after rinsing 3 × 30 min with 1% PBST at 25°C, samples were stored in 1× PBS at 4°C until imaged. Samples were imaged using one of the following confocal microscopes: an inverted Zeiss LSM 880 confocal using either the 10× EC-Plan NeoFluar (NA = 0.3, working distance = 5.2 mm), or 25× multi-immersion LD LCI Plan-Apochromat (NA = 0.8, working distance = 0.57 mm), an upright Zeiss LSM 800 confocal using 20× W immersion Plan-Apochromat (NA = 1.0, working distance = 2.4 mm), or a Leica DM6 CFS STELLARIS upright confocal (Leica Microsystems) with LAS X software (V4.1.1.23273) using HC APO 10×/0.3 W U-V-I (NA = 0.3, working distance = 3.6 mm), HC FLUOTAR L 25×/0.95 VISIR (NA = 0.95, working distance = 2.4 mm), and HC APO L 63×/0.9 W U-V-I CS2 (NA = 0.9, working distance = 2.2 mm). Z-stack and tile functions were used to capture the entirety of the biceps and triceps muscles. Widefield images were acquired using a Leica M80 stereo microscope.

### Forelimb optical clearing

To visualize the 3D morphology and spatial patterning of muscle cells embedded within the developing limb, and confirm that ECM structure was maintained after decellularization, SeeDB, a fructose-based clearing solution, was utilized. Forelimbs were cleared following (Calve et al., 2015; Ke et al., 2013). Fructose solutions of varying concentrations (20%, 40%, 60%, 80%, 100%, and 115% wt/vol) were generated by dissolving D-(-)-fructose (JT Baker) in Milli-Q water with 0.5% α-thioglycerol (Sigma-Aldrich) to prevent browning and 0.02% sodium azide to prevent fungal infection in the higher concentration solutions. Fixed and stained tissues were equilibrated to increasing concentrations of fructose by incubating in each formulation for 8-24 h under gentle rocking at RT.

### Mechanical testing of limbs

Limbs were collected and decellularized as described in the forelimb decellularization section but not fixed. After blocking, the limbs were stained with AF488-conjugated WGA and an antibody to COL22A1 as described above. Methods were modified from (Acuna et al., 2021). In brief, forelimbs were held using two bulldog clamps (Fine Science Tools) attached to the wrist and shoulder respectively, taking care to avoid the long triceps tendon. Clamps were secured in a custom uniaxial loading rig controlled by a microrobotic system (FT-RS1002, FemtoTools) with samples submerged in PBS, and the humerus was cut from the bicep side to allow ECM fibers in the triceps to stretch freely during testing. The loading system was integrated under a confocal microscope for imaging at 25×. To evaluate deformation in the ECM, rectangles were photobleached on either side of the MTJ to be used as fiducial markers. Nine laser lines centered at 488 nm were set to 100% power, and a z-stack was acquired to photobleach the markers. After photobleaching, a reference z-stack was taken of the tendon, MTJ, and muscle area. The distal end of the forelimbs was then displaced 2.5 mm to stretch the limb, and a second z-stack was acquired for the same region of interest. Forelimbs were returned to the reference configuration to generate a third z-stack for the region of interest.

### Image processing

Confocal stacks were adjusted for brightness and contrast then rendered in 3D using FIJI (NIH). For vibratome sections, image stacks were processed into z-projections using FIJI. For mechanical testing, image stacks were processed into summed z-projections using FIJI. To remove some of the optical distortion inherent to confocal microscopy, the Nearest Neighbor Deconvolution function within the ZenBlue software (Carl Zeiss Microscopy) or LAS X software (Leica Microsystems) was applied. Figures were assembled by taking snapshots of 3D-rendered image volumes or selecting 2D slices from stacks and were arranged using Adobe Photoshop and Illustrator.

### Proteomic Analysis

Forelimbs were microdissected from E13.5 *Pax3*^*Cre/Cre*^ or WT embryos, the autopod region was removed, and the remaining tissue was snap frozen and stored at −80°C. Dissected limbs from one embryo were pooled and processed for mass spectrometry as described in (Jacobson et al., 2020b), with slight modifications. Briefly, proteins were fractionated into five samples (C, N, M, CS, IN) using homemade buffers (Jacobson et al., 2020a), enzymatically digested into peptides, and peptides were analyzed by liquid chromatography-tandem mass spectrometry (LC-MS/MS).

Digested peptides were analyzed in an Dionex UltiMate 3000 RSLC nano System (Thermo Fisher Scientific) coupled on-line to Orbitrap Fusion Lumos Mass Spectrometer (Thermo Fisher Scientific) as previously described (Barabas et al., 2019). Briefly, reverse phase peptide separation was accomplished using a trap column (300 μm ID × 5 mm) packed with 5 μm 100 Å PepMap C18 medium coupled to a 50-cm long × 75 µm inner diameter analytical column packed with 2 µm 100 Å PepMap C18 silica (Thermo Fisher Scientific). The column temperature was maintained at 50°C. One µg peptides (equivalent volume) was loaded to the trap column in a loading buffer (3% acetonitrile, 0.1% FA) at a flow rate of 5 µL/min for 5 min and eluted from the analytical column at a flow rate of 200 nL/min using a 130-min LC gradient. Column was washed and equilibrated by using three 30-min LC gradient before injecting next sample. All data were acquired in the Orbitrap mass analyzer and data were collected using an HCD fragmentation scheme. For MS scans, the scan range was from 350 to 1600 *m*/*z* at a resolution of 120,000, the automatic gain control target was set at 4 × 10^5^, maximum injection time 50 ms for both MS1 and MS2, and dynamic exclusion was 30s. MS data were acquired in Data Dependent mode with cycle time of 5s/scan. MS/MS data were collected at a resolution of 7,500 with isolation window of 1.2 m/z. The RF lens and collision energy were set at 30%.

Raw data files were processed by MaxQuant (Cox and Mann, 2008), and raw or LFQ intensities were used for downstream data analysis (Cox et al., 2014). Identified proteins were annotated for Cellular Compartment and Matrisome classifications (Naba et al., 2012; Saleh et al., 2019), and the distribution of these classifications were calculated as a percentage of total raw intensity (Jacobson et al., 2020b). For volcano plot analysis of ECM proteins identified in the IN fraction, LFQ intensities were log_2_ transformed, the fold-change was calculated between Pax3^*Cre/Cre*^ and WT samples, and a two-tailed Student’s t-test was used for statistical comparisons. LFQ intensities were also used for heatmap analyses, wherein proteins were first clustered by Matrisome classification and then by abundance. Raw intensities were used for ECM protein identification in the other fractions.

## Data availability statement

The data supporting the findings of this study are openly available in the MassIVE repository at MSV000088620 and will be released upon publication. *Reviewers: please see Supplemental Information to find how to access the data*.

## Author contributions

Conceptualization: S.C. Methodology: Y.L., S.N.L., H.L., K.R.J; Validation: S.N.L.,Y.L., Y.B., H.L.; Formal analysis: Y.L., S.N.L., H.L., K.R.J.; Investigation: Y.L, S.N.L, Y.B., H.L., K.R.J.; Data curation: S.N.L., Y.B., H.L., K.R.J.; Writing – original draft: Y.L., S.N.L., S.C.; Writing - review & editing: Y.L., S.N.L., H.L., K.R.J., S.C.; Visualization: Y.L., S.N.L, Y.B., H.L., K.R.J.; Supervision: S.C.; Project administration: S.C.; Funding acquisition: S.C.

## Acknowledgments

We would like to thank members of the Calve lab for helpful discussions, notably Julian M. Jimenez and Callan Luetkemeyer, PhD for mechanical instrumentation development and Dalton Miles for MTJ integrity discussions. We also would like to thank Karin (Kaisa) Ejendal, PhD for assistance with mice.

## Grants

This work was supported by the National Institutes of Health [DP2 AT009833 and R01 AR071359 to S.C.] and the Bilsland Dissertation Fellowship [to Y.L.].

## Notes

### Competing Interest Statement

The authors have declared no competing interest.

